# circHiC: circular visualization of Hi-C data and integration of genomic data

**DOI:** 10.1101/2020.08.13.249110

**Authors:** Ivan Junier, Nelle Varoquaux

## Abstract

Genome wide contact frequencies obtained using Hi-C-like experiments have raised novel challenges in terms of visualization and rationalization of chromosome structuring phenomena. In bacteria, display of Hi-C data should be congruent with the circularity of chromosomes. However, standard representations under the form of square matrices or horizontal bands are not adapted to periodic conditions as those imposed by (most) bacterial chromosomes. Here, we fill this gap and propose a Python library, built upon the widely used Matplotlib library, to display Hi-C data in circular strips, together with the possibility to overlay genomic data. The proposed tools are light and fast, aiming to facilitate the exploration and understanding of bacterial chromosome structuring data. The library further includes the possibility to handle linear chromosomes, providing a fresh way to display and explore eukaryotic data.

**Availability and implementation:** The package runs under Python 3 and is freely available at https://github.com/TrEE-TIMC/circHiC. The documentation can be found at https://tree-timc.github.io/circhic/; images obtained in different organisms are provided in the gallery section and are accompanied with codes.

## I. INTRODUCTION

Genome wide high-throughput chromosome conformation capture (Hi-C) methods (Lieberman-Aiden *et al*. 2009) and related techniques (Denker and De Laat 2016) allow to compute, up to an unknown global factor, contact frequencies between any two loci along a genome. Rationalization of chromosome structuring phenomena requires, in the first place, an appropriate visualization of these data. Several challenges along this line have been raised and (sometimes) solved. For instance, eukaryotic data can be browsed at different scales, under the form of square matrices (Durand *et al.* 2016, Kerpedjiev *et al*. 2018, Yardimci and Noble 2017).

Compared to eukaryotes, bacterial chromosomes are much smaller, yet they also harbor a complex multilayer organization (Lagomarsino *et al*. 2015). Most importantly, most bacterial chromosomes are circular. From the perspective of Hi-C visualization, this raises the ineluctable problem of representing a physical system with spherical symmetry using a two-dimensional Euclidean space. That is, the standard square matrix or horizontal band representations commonly used for Hi-C data are not appropriate for a system with periodic conditions as those associated with bacterial chromosomes. In particular, there is currently no tool, to the best of our knowledge, allowing to display Hi-C heat maps in a circular strip.

Here, we fill this gap by developing a Python library, `circhic`, that efficiently displays bacterial Hi-C data accordingly. The library further includes the possibility to overlay genomic data. It also includes the possibility to handle linear chromosomes and, hence, can be used to visualize eukaryotic data.

## II. ALGORITHM

Let *H* be the input Hi-C matrix (size *N × N*). Our algorithm consists in using a system of polar coordinates to project *H* onto a circular strip (Figure 1). Specifically, a circle of the strip corresponds to all contact frequencies between pairs of loci that are separated by a certain genomic distance, *s*. In this context, `circhic` first consists in specifying the genomic distances *s*_*in*_ and *s*_*out*_ for the inner and outer circles – we note here that `circhic` displays by default, in the inward and outward parts of the strip, contacts corresponding to *s ∈* [0, *s*_*in*_] and to *s ∈* [0, *s*_*out*_], respectively. We then consider, in each of these two parts of the strip, a linear relationship between the radius of a circle, *r*_*s*_, and the corresponding genomic distance, *s*. Altogether, we eventually have 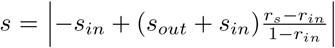 for the entire strip, with *r*_*in*_ the radius of the inner circle and where all the radius are normalized by the radius of the outer circle (i.e. *r*_*out*_ = 1). Next, any pair of loci can be written as 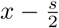 and 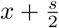, with *x* and *s* the respective associated middle locus and genomic distance – note that loci (i.e. indexes of *H*) are considered modulo *N*. The associated contact frequency is then properly positioned along its associated circle (with radius *r*_*s*_) by specifying the polar angle of *x*. To this end, we use a clockwise, twelve o’clock origin angle *θ*_*x*_ that is linear in 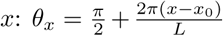. *x*_0_, which can be set by the user, corresponds to the middle locus of the pairs of loci whose contact frequencies are displayed vertically.

**Fig. 1.**
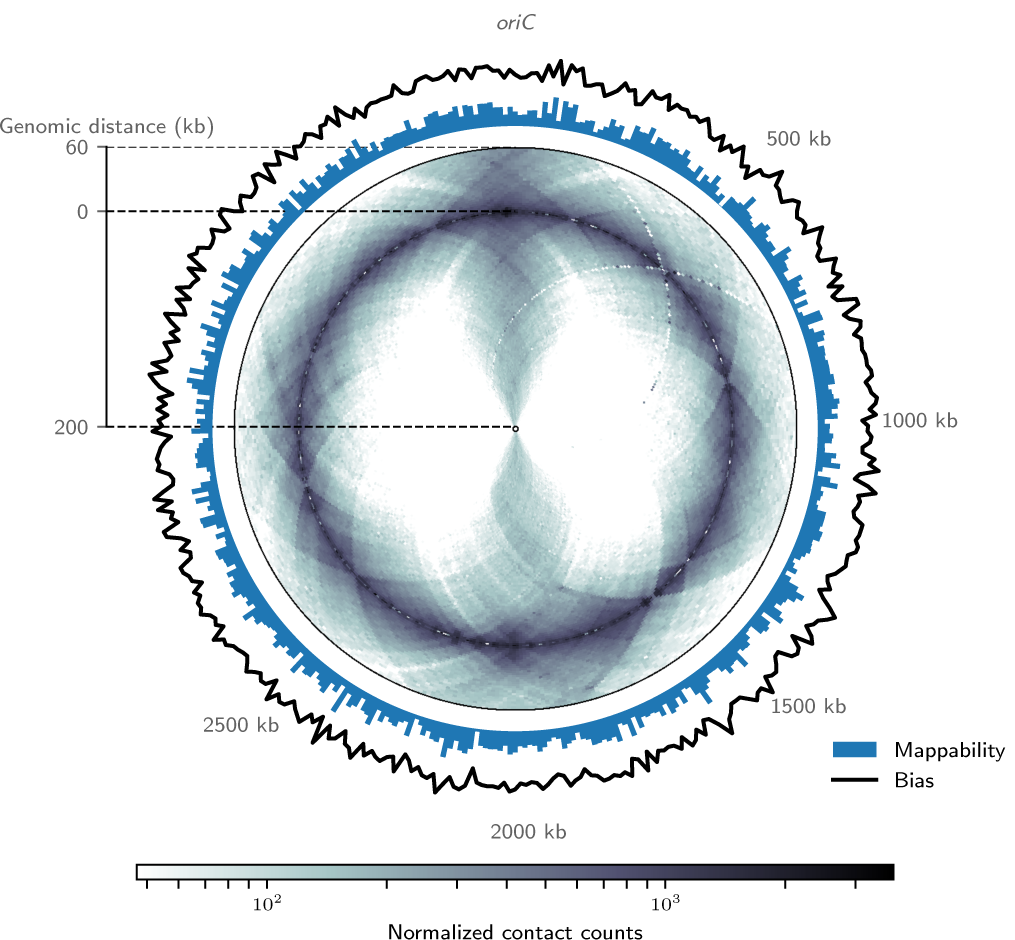
Circular visualization of Hi-C data obtained in *C. crescentus* (Le *et al*. 2013) *generated using circhic in a default ‘reflect’ mode, using s*_*in*_ = 200 kb and *s*_*out*_ = 60 kb. Two additional genomic data (the bias associated with the ICE normalization of data (Imakaev *et al*. 2012) in black lines and cumulative raw contact counts in black bars) are displayed outside the Hi-C data.

Using this framework, `circhic` consists in filling up the entries of a matrix *C* (size *N*_*c*_ *× N*_*c*_) that are located in the circular strip centered in 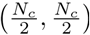 with inner radius 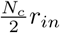 and outer radius 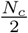, by considering the corresponding entries of *H*. To this end, we consider polar coordinates associated with the circular strip and we use, in place of the trigonometric angle, the above clockwise angle with twelve o’clock origin. Each entry (*i*_*c*_, *j*_*c*_) of the circular strip in *C* is thus associated with a coordinate (*r, θ*) with 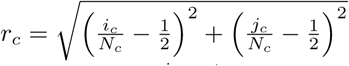 and *θ*_*c*_ such that 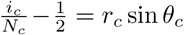 sin *θ*_*c*_ and 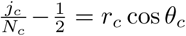. Given (*r*_*c*_, *θ*_*c*_) and following the discussion of the previous paragraph, we then set 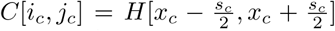 with 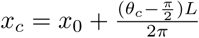 and 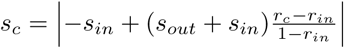.

Note that, just as with square displays, data on each side of *s* = 0 are redundant (here up to min(*s*_*in*_, *s*_*out*_)), generating a reflection effect. This allows for instance to enhance the presence of chromosomal interaction domains as in the case of *Caulobacter crescentus* (Le *et al*. 2013) (Figure 1). In addition to this ‘reflect’ mode, `circhic` provides a ‘distant’ mode to highlight contacts away from the main diagonal (see gallery in the online documentation). Note finally that *N*_*c*_ is larger or equal to *N*. We thus define the granularity of the circular projection by *N/N*_*c*_, equal to 0.5 by default. The smaller the granularity is, the neater the circular display, at the cost of a longer computation.

## III. IMPLEMENTATION

We built upon the popular visualization library Matplotlib (Hunter 2007), using core libraries of the scientific Python ecosystem (`numpy`, `pandas`, …). `circhic` relies on an object oriented approach, where a figure (the core object of the library) holds all the necessary elements for the visualization: the size of the genome, the orientation of the plot, and a number of information relating to the size and shape of the figure. A number of methods can then be executed to visualize the data: `plot hic` provides support for transforming the square contact count matrix into a circular one, and `plot lines`, `plot bars`, … allow to overlay relevant genomic information onto the contact count matrix.

## IV. RESULTS

As an example, in Figure 1 we show a circular representation of the first high-resolution Hi-C data reported in a bacterial chromosome (Le *et al.* 2013) together with genomic data. In the gallery section of the online documentation, we provide additional examples using publicly available data, including the visualization of a linear human chromosome.

## V. CONCLUSION

`circhic` provides a useful, fast representation of Hi-C data that respects the circularity of bacterial chromosomes. The possibility to overlay genomic information aims at facilitating the exploration and understanding of chromosome structuring data. In practice, the build upon Matplotlib allows a great flexibility to generate complex figures.

